# RL-Finetuning of OpenAI o1-mini to Enhance Biomedical Reasoning

**DOI:** 10.1101/2025.05.19.654988

**Authors:** Kyle Swanson, Yiqun T. Chen, Aaron Jaech, James Zou

**Affiliations:** Department of Computer Science, Stanford University; Departments of Biostatistics and Computer Science, Johns Hopkins University; Department of Biomedical Data Science, Stanford University; OpenAI

**Author notes:** Equal contribution (Kyle Swanson, Yiqun T. Chen).

## Abstract

Recent breakthroughs in advanced reasoning large language models (LLMs), such as OpenAI’s o1, have achieved impressive results in domains like math and coding. However, it’s not clear how much this type of reasoning helps in solving biomedical problems that involve more domain specialized knowledge and open-ended reasoning. Across two biomedical domains—gene characterization and small molecule property prediction—we find that the commercially available o1-mini model does not consistently outperform non-reasoning LLMs like GPT-4o. This motivated us to explore how much we can improve o1-mini’s biomedical reasoning through reinforcement learning (RL) finetuning. We show that RL finetuning of o1-mini results in large improvements in performance on gene classification, where it surprisingly outperformed domain-specific state-of-the-art models on some tasks. The results are mixed for small molecule prediction, suggesting that chemical reasoning could be more challenging for LLMs. We conclude with a discussion of the challenges and takeaways from this initial exploration of RL finetuning reasoning models for biomedical tasks.

## Introduction

Recent advances in large language models (LLMs) have demonstrated impressive success in natural language understanding and generation, marking remarkable linguistic fluency and knowledge retention. However, these models often struggle with tasks that require systematic reasoning, e.g., when logical inference and problem-solving with domain knowledge are involved. While LLMs can generate linguistically coherent and plausible answers, the answers often exhibit reasoning inconsistencies and errors, limiting their reliability and usability in applications such as scientific discovery.

To bridge this gap, recent research on LLMs has explored various methodologies, including Chain-of-Thought (CoT) (Wei et al. 2022) and TextGrad (Yuksekgonul et al. 2025), as well as finetuning with reinforcement learning to enhance reasoning consistency. These approaches aim to improve models’ reasoning abilities. Notably, models such as OpenAI’s o1 and o3-mini exemplify the latter approach—they are LLMs trained with reinforcement learning to perform multi-step, complex reasoning. At a high level, these models generate an extended internal chain of thought (also known as reasoning tokens) before responding to user prompts. This training methodology prioritizes accurate, multi-step reasoning over one-shot responses, which is common in non-reasoning LLMs. Consequently, reasoning models have demonstrated impressive performance in domains requiring complex problem-solving, such as coding and math.

This observation has also motivated researchers to explore whether advances in reasoning models can be applied to challenges at the intersection of biomedicine and AI, where many tasks are reasoning-intensive and push the limits of out-of-the-box LLMs. For example, understanding the role of a gene in a genetic pathway requires biologists first to recall the functionalities of the genes involved and then infer their roles based on experimental data from the literature. Similarly, predicting the properties of a chemical compound (such as toxicity or solubility) from its structure or description demands reasoning through known chemical substructures and rules. Even very large language models may struggle with these tasks because they lack structured reasoning in fundamental biology and chemistry, often relying on popular associations present in their pretraining data.

On the other hand, biomedical AI has long leveraged domain-specific models optimized for specific tasks. Graph neural networks (GNNs) excel at molecular property prediction by representing chemical structures as graphs, effectively capturing atomic interactions (Heid et al. 2024). Gene embedding models like Geneformer (Theodoris et al. 2023) and GenePT (Chen and Zou 2024) specialize in gene function prediction using transformer-based architectures trained on biological data. Meanwhile, biomedical BERT variants (e.g., BioBERT (Lee et al. 2020) and PubMedBERT (Gu et al. 2022)) perform well in text-based biomedical NLP tasks such as entity recognition and literature classification. While these models achieve high accuracy in specialized domains, they lack flexibility compared to LLMs: GNNs cannot process unstructured text, Geneformer is limited to genomic applications, and small-scale language models such as BioBERT struggle with multi-step reasoning.

To address this gap, we investigate whether finetuning a reasoning LLM can integrate specialized biomedical knowledge while preserving its multi-step reasoning capabilities. By adapting OpenAI’s o1-mini, we evaluate whether a finetuned reasoning LLM can rival domain-specific models across two key tasks:

- **Gene classification** – Determining the functional category of a gene or variant in different contexts.
- **Small molecule property prediction** – Predicting properties of a chemical compound (e.g., toxicity, solubility, or antibiotic activity) based on its structure or description.

Our results show that reinforcement learning (RL) finetuning of o1-mini significantly improves the model’s performance in gene classification. However, outcomes for small molecule property prediction are mixed. Additionally, we found that class rebalancing helps mitigate data imbalances, further boosting finetuned performance across different tasks.

Overall, our findings highlight the potential of biomedically adapted reasoning models to outperform both standard LLMs and specialized approaches in complex problem-solving.

## Results

We investigate the impact of finetuning LLMs on two classes of biological problems: gene classification and small molecule property prediction.

### Models

In all experiments, we evaluate both pre-trained and finetuned versions of OpenAI’s o1-mini and GPT-4o-mini (GPT-4o for gene classification), referred to as “o1-base” and “4o-base” for the pre-trained models and “o1-finetuned” and “4o-finetuned” for our RL finetuned versions. We finetune o1-base using reinforcement learning (RL) and 4o-base using supervised finetuning, both provided by OpenAI (see Methods).

We compare these LLMs against two types of baseline models. First, we establish simple statistical baselines. For classification accuracy, we use a majority class baseline, which measures accuracy when always predicting the most common answer in the test set. For F1 scores, we use an “all ones” baseline, representing the F1 score when all predictions are ones.

Second, we compare the LLMs to domain-specific machine learning models. For gene classification, we benchmark against GenePT, an embedding model derived from textual descriptions of the functionalities of genes and cells that have competitive performance in gene classification tasks. In each of these cases, we train a regularized logistic regression model on the GenePT embedding features and generate predictions on the test set. For small molecule datasets, we train a domain-specific graph neural network, Chemprop-RDKit (Swanson et al. 2024b), on the same training, validation, and test sets as the LLMs.

### Data

We selected a number of datasets across two categories of biological questions: gene functionality classification and small molecule property prediction.

#### Gene Functionality Classification

To probe language models’ understanding of gene functionality, we used the following datasets used in the analysis in Theodoris et al. (2023) and Chen and Zou (2024).

1. **Dosage Sensitivity:** A binary classification problem distinguishing dosage-sensitive versus dosage-insensitive transcription factors (TFs). There are 487 data points after preprocessing (122 sensitive, 365 insensitive). Finetuning for o1-mini was performed on a randomly selected 50% training dataset, and metrics for all models are reported on the other 50% held-out dataset.
2. **Promoter Classification:** A three-class classification problem categorizing promoter genes into bivalent (N=106), Lys4-only-methylated (N=78), or non-methylated (N=41). Finetuning for o1-mini was performed on a randomly selected 50% training dataset, and metrics for all models are reported on the other 50% held-out dataset.
3. **Genome-wide Promoter Classification:** Genome-wide profiling of bivalent (N=2,602) versus Lys4-only-methylated (N=7,213) promoter domains. Finetuning for o1-mini was performed using a class-balanced training set (subsampled 50% of the data to achieve a 1:1 ratio of bivalent versus Lys4-only-methylated examples). Metrics for all models are reported on the other 50% held-out dataset without class-balancing.
4. **Long vs. Short-Range TFs:** A binary classification problem distinguishing long-range (N=46) versus short-range (N=130) TFs. Finetuning for o1-mini was performed using a class-balanced training set (subsampled 50% of the data to achieve a 1:1 ratio of short-versus long-range examples). Metrics for all models are reported on the other 50% held-out dataset without class-balancing.
5. **NOTCH1 Targets:** A binary classification problem identifying NOTCH1 (N1)-activated genes (N=563) versus non-targets (N=563) in the gene network governing cardiac valve disease, as mapped by Theodoris et al. (2015, 2021). Finetuning for o1-mini was performed on a randomly selected 50% training dataset, and metrics for all models are reported on the other 50% held-out dataset.
6. **NOTCH1 Centrality:** A binary classification problem identifying genes that are central (N=98) versus peripheral (N=192) in the NOTCH1 (N1) network governing cardiac valve disease, as mapped by Theodoris et al. (2015, 2021). Finetuning for o1-mini was performed using a class-balanced training set (subsampled 50% of the data to achieve a 1:1 ratio of central versus peripheral examples). Metrics for all models are reported on the other 50% held-out dataset without class-balancing.

#### Small Molecule Property Prediction

For small molecule property prediction, we use six binary classification datasets from the Therapeutics Data Commons (TDC) ADMET benchmark (Huang et al. 2021). This benchmark contains datasets of small molecules with measurements of their absorption, distribution, metabolism, excretion, and toxicity (ADMET) properties. Determining these properties is crucial for developing safe and effective small molecule drugs.

1. **AMES:** AMES is a bacterial test of mutagenicity, which is the ability of a drug to induce genetic alterations that can lead to cell death or other adverse effects.
2. **Bioavailability:** Oral bioavailability is the rate and extent to which a drug is absorbed and becomes available at the desired site of action.
3. **CYP2D6 Inhibition:** This assay measures the amount that a drug inhibits CYP2D6, which is an enzyme involved in the metabolism of various molecules.
4. **CYP3A4 Substrate:** This assay measures the amount that a drug is metabolized by CYP3A4, which is another metabolic enzyme like CYP2D6.
5. **CYP3A4 Inhibition:** This assay measures the amount that a drug inhibits CYP3A4.
6. **P-gp:** P-glycoprotein (P-gp) is a protein involved in intestinal absorption, drug metabolism, and brain penetration. This assay measures how much a drug inhibits P-gp, which can alter the drug’s bioavailability and safety.

For each dataset, we use the train, validation, and test splits from the TDC using scaffold splits with seed 1. If any of the train, validation, or test splits have more than 2,000 molecules, we randomly sample 2,000 molecules from that split.

We additionally explored a small-molecule antibiotics dataset (Swanson et al. 2024a), but we defer those details to the Appendix since they are qualitatively similar to the TDC dataset results.

### Experiments

#### Gene Classification

Across various gene classification tasks, finetuned LLM models (4o-finetuned and o1-finetuned) generally outperform their base counterparts, with particularly strong improvements in the dosage sensitivity classification task (Figure 1, Supplementary Figure 1). In particular, the RL finetuned o1-finetuned model demonstrates the best prediction performance across tasks (0.75 accuracy; 0.75 F1), followed closely by GenePT (0.74 accuracy; 0.72 F1). Additionally, finetuning for o1 generally yields a larger improvement over the base model compared to 4o, with an average increase in accuracy of 8% for o1 models versus less than 1% for 4o models. This difference is partly attributable to the high variance in 4o finetuning performance, where we observe substantial improvement for tasks like dosage sensitivity but decreased accuracy for promoter classification.

**Figure 1:**
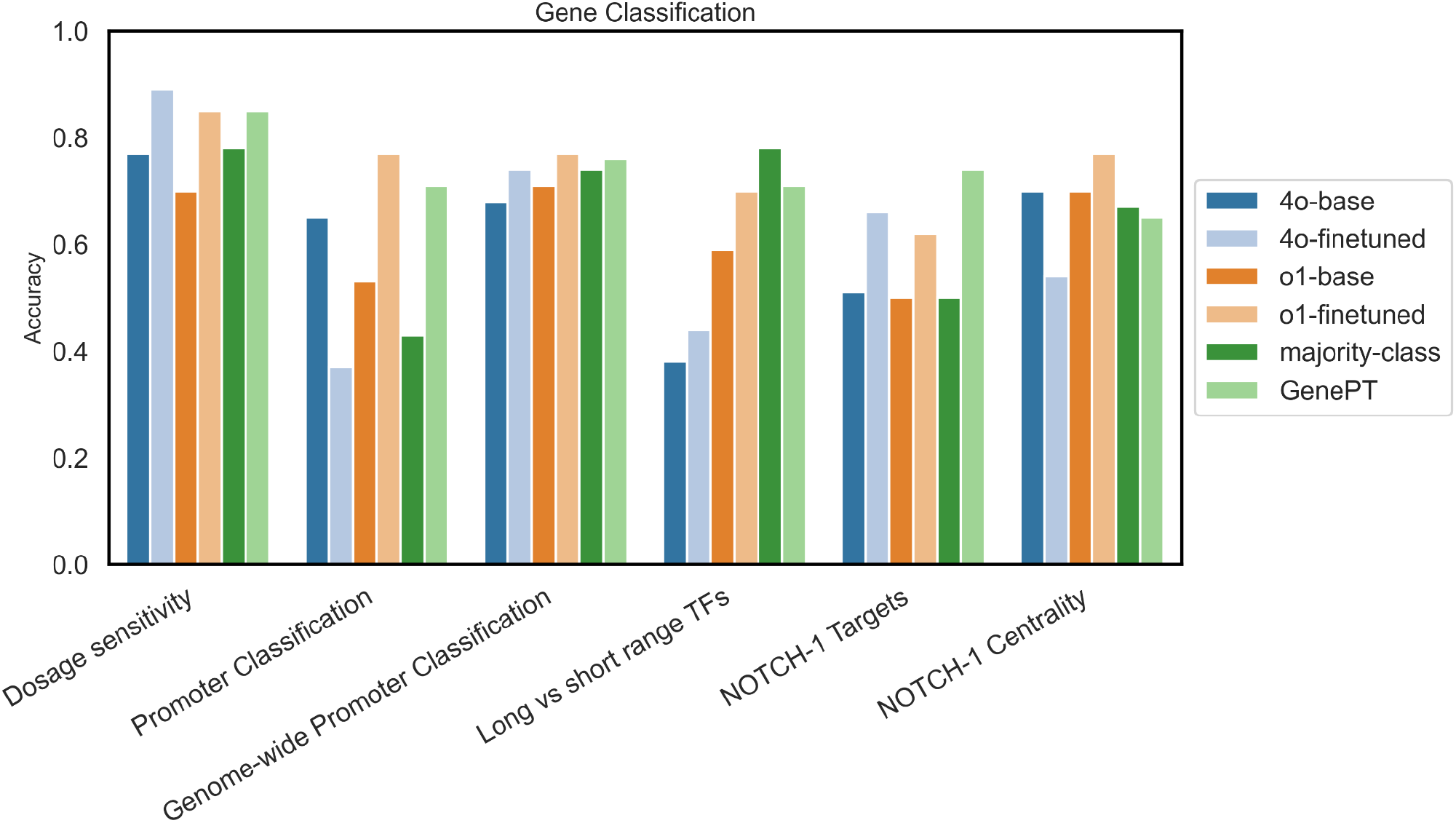
Accuracy scores comparing LLM models (before and after finetuning), a majority class baseline, and logistic regression applied to GenePT embeddings on the test sets of gene classification datasets. o1-finetuned has the highest average accuracy across tasks (0.75), followed by GenePT (0.74) and majority class (0.65).

Fine-tuning also altered the reasoning patterns of the o1-mini model. Below, we present three randomly sampled examples from different tasks. Due to OpenAI policies, we can only display the summarized reasoning rather than the full reasoning trajectory.

##### o1-base reasoning

COMTD1 is associated with active histone mark H3K4me3 without repressive marks, indicating a Lys4-only state. Prediction: Lys4-only. Correct answer: non-methylated.

##### o1-finetuned reasoning

COMTD1 has a non-methylated promoter typically associated with housekeeping genes. Prediction: non-methylated. Correct answer: non-methylated.

##### o1-base reasoning

ZNF527 is a transcription factor whose dosage alterations can impact gene regulation and cellular functions, indicating dosage sensitivity. Prediction: dosage sensitive. Correct answer: dosage insensitive

##### o1-finetuned reasoning

ZNF527 is a zinc finger protein transcription factor. Due to the lack of known associations with dosage sensitivity from available genomic knowledge, it is classified as dosage-insensitive. Prediction: dosage insensitive. Correct answer: dosage insensitive

##### o1-base reasoning

NCOR1 acts as a corepressor influencing gene expression indirectly within the NOTCH1-dependent network governing cardiac valve disease. Prediction: N1-peripheral. Correct answer: N1-central.

##### o1-finetuned reasoning

NCOR1 is a nuclear receptor corepressor that interacts with transcriptional regulators, playing a crucial role in repressing gene transcription. Given its function in gene regulation and likely involvement in pathways downstream of NOTCH1, it is classified as central within the NOTCH1-dependent gene network. Prediction: N1-central. Correct answer: N1-central.

Before finetuning, the o1-base model tends to give generic biological associations and make overly simplistic inferences, such as assuming all transcription factors are dosage sensitive or incorrectly classifying promoter states based on partial histone mark information. After finetuning, the o1-finetuned model demonstrates slightly more nuanced domain knowledge, incorporating specific functional genomic concepts and making more precise inferences based on molecular function. This improved reasoning leads to more accurate predictions, as seen in the COMTD1 promoter classification, where the model correctly identifies non-methylated status rather than relying solely on histone marks, and in the ZNF527 case, where it properly recognizes that zinc finger proteins are not inherently dosage sensitive.

#### Small Molecule Property Prediction

Overall, finetuning LLMs—both o1-finetuned and 4o-finetuned—on the TDC ADMET datasets improves accuracy (Figure 2) and F1 scores (Supplementary Figure 2). However, these improvements can often be attributed to the model simply learning to predict the majority class (active or inactive) for most if not all molecules rather than the model learning to truly differentiate between active and inactive molecules. Reflecting this result, the LLMs are rarely able to outperform a simple majority class baseline or the domain-specific Chemprop GNN trained on the same data.

**Figure 2:**
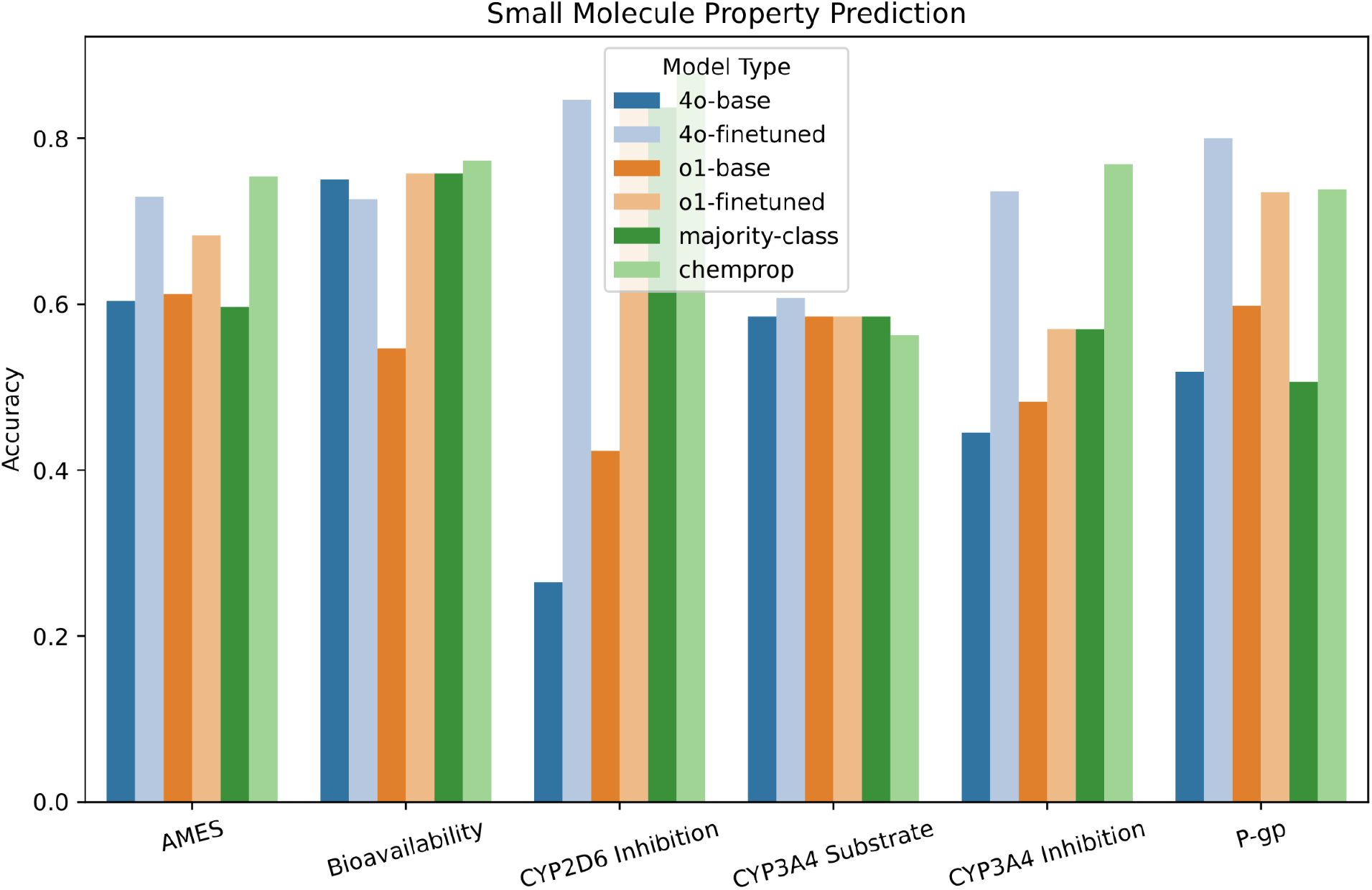
Accuracy scores comparing LLM models (before and after finetuning), a majority class baseline, and a Chemprop GNN model on the test sets of the TDC ADMET datasets.

##### AMES

Finetuning improves upon both the 4o-base and o1-base models and they outperform the majority class baseline, but they cannot match the performance of the Chemprop model. Furthermore, even though o1-finetuned performance improves, the explanations become less informative. For example, for one molecule, the explanation of the o1-base model is “The molecule O=Cc1ccc(CO)o1 does not contain structural features commonly associated with mutagenicity, such as nitro groups or aromatic amines. Its structure lacks known mutagenic functional groups, suggesting it is likely inactive in inducing genetic alterations.” However, the explanation of the o1-finetuned model is simply “The molecule does not contain known mutagenic groups.”

##### Bioavailability

No model can significantly beat the majority baseline. Interestingly, finetuning 4o hurts performance while finetuning the o1-base model improves performance. However, o1-finetuned collapses to predicting active for every molecule (the true distribution is 76% active).

##### CYP2D6 Inhibition

Finetuning improves LLM model performance but only because the models collapse to nearly always (4o-finetuned) or always (o1-finetuned) predicting inactive (the true distribution is 84% inactive). Chemprop outperforms all of the LLMs.

##### CYP3A4 Substrate

The LLM models outperform Chemprop, but they are still only on par with the majority class baseline. The o1-finetuned model collapses to predicting all active molecules (the true distribution is 59% active).

##### CYP3A4 Inhibition

Finetuning improves LLM performance and 4o-finetuned even outperforms the majority class baseline. However, o1-finetuned collapses to predicting all inactive molecules (the true distribution is 57% inactive), and Chemprop outperforms all of the LLMs.

##### P-gp

The finetuned LLMs outperform both Chemprop and the majority class baseline. Although o1-base outperforms 4o-base, 4o-finetuned outperforms o1-finetuned.

## Discussion

Finetuning the o1-mini reasoning model for biomedical tasks shows promise, but the benefit of finetuning depends on the particular task and dataset. On the gene classification tasks, o1-finetuned significantly outperformed o1-base as well as 4o-finetuned and often matched or exceeded the performance of a domain-specific model, GenePT. However, the results on the small molecule property prediction datasets were markedly different. While finetuning often significantly improved the performance metrics, this was typically via a collapse of the model to always predicting the majority class, which may have high accuracy but has no useful predictive value. Therefore, the efficacy of finetuning reasoning models like o1-mini may depend on the specific biomedical domain (genes or small molecules) since those domains have different data representations (gene names or small molecule SMILES) and different dataset characteristics (size, class balance, etc.).

One notable limitation of RL finetuning o1-mini is its sensitivity to class imbalance. In four of the six small molecule property prediction datasets, o1-finetuned collapses to predicting the majority class on every test molecule. While this might be reasonable for very imbalanced datasets like CYP2D6 Inhibition with 84% inactive molecules, it also occurs for even mildly imbalanced datasets like CYP3A4 Inhibition with 57% inactive molecules. This has also been observed in gene classification tasks where class imbalance is present (e.g., Genome-wide Promoter Classification and NOTCH1 Centrality), where finetuning on a class-balanced dataset achieved better results, even though the final target distribution remained imbalanced. This indicates that RL finetuning may easily fall into short-sighted solutions like predicting the majority class at the cost of identifying more advanced reasoning traces that can truly analyze the properties of the small molecules.

Another challenge of RL finetuning o1-mini is that it is challenging to determine how the reasoning of the model is changing to improve performance. The internal reasoning of the model is not available to users, so only post-hoc explanations provided alongside the answer are available to analyze the model’s reasoning. When finetuning leads the model to collapse to predicting the majority class, these explanations often collapse as well to an empty string. Even when finetuning is successful and improves test set performance, the post-hoc explanations often do not provide much information, as illustrated by the reduction in explanation complexity in the AMES explanations despite performance improvement. Thus, it remains unclear whether RL finetuning is actually causing the model to engage in more complex reasoning or whether the model is simplifying identifying shortcut solutions like majority class prediction.

Despite these limitations, the results presented here indicate that there is promise for RL finetuning of reasoning models for biomedical tasks. While we explore the o1-mini model here as a proof of concept, newer reasoning models such as o3-mini (Zhang et al. n.d.) may yield better results. Future work should further explore the task and dataset conditions that are required for successful finetuning of reasoning models in the biomedical domain. This will allow researchers to determine when to finetune a reasoning model for biomedical tasks and when to use either the base LLM or a domain-specific tool.

## Methods

### Finetuning

We start with the pre-trained base models o1-mini-2024-09-12 (“o1-base”) and either gpt-4o-2024-08-06 for gene classification or gpt-4o-mini-2024-07-18 for small molecule property prediction (“4o-base”) and finetune them using OpenAI APIs to create models we refer to as “o1-finetuned” and “4o-finetuned,” with a distinct finetuned model for each dataset. During finetuning, both model types are instructed to provide their answers in JSON format, and we use all default hyperparameters unless otherwise specified.

During RL finetuning of the o1-base model, the LLM-generated answer is extracted from the LLM-generated JSON object and is compared to a reference answer using a grader to compute a reward that is used to finetune the model. We used the `string-check-grader` with the operation `equals`, which gives a reward of 1 if the answer provided by the model exactly matches the reference answer and a reward of 0 otherwise. The LLM-generated explanation does not affect the finetuning reward and is only used for human interpretation of the LLM-generated answers.

During supervised finetuning of the 4o-base model, the LLM-generated JSON string is directly compared to a reference JSON string, which contains the reference answer and an empty string for the explanation. Therefore, the 4o-base model produces explanations but the 4o-finetuned models generate empty strings for explanations. While this may not be ideal from an interpretability standpoint, we did this so that both o1-style and 4o-style models use identical prompts and nearly identical output formats during finetuning, with only the model architecture and finetuning procedure differing.

During evaluation, any invalid answers (e.g., invalid JSON) are ignored and not included in the evaluation. Note that this may result in slightly different test sets for each model.

### Gene Classification Prompts

#### Dosage Sensitivity

~~~
Use biological knowledge on human genomics to answer the following
binary classification question in the context of whether a
transcriptional factor is dosage sensitive or dosage insensitive.
Answer in one of the two categories: sensitive or insensitive and
respond with a JSON object where each element contains ‘gene’,
‘answer’, and a short ‘explanation’ without any other supporting
commentary or formatting. Gene name: <Gene name>
~~~

#### Promoter Classification

~~~
Use biological knowledge on human genomics to answer the following
classification question that classifies human promoter genes into
different classes. Answer with one of these categories: bivalent,
Lys4-only, and non-methylated and respond with a JSON object
containing ‘gene’, ‘answer’, and a short ‘explanation’ without any
other supporting commentary or formatting. Gene name: <Gene name>
~~~

#### Genome-wide Promoter Classification

~~~
Use biological knowledge on human genomics to answer the following
classification question that classifies transcriptional factors into
different classes. Answer with one of these categories: bivalent,
Lys4-only and respond with a JSON object containing ‘gene’, ‘answer’,
and a short ‘explanation’ without any other supporting commentary or
formatting. Gene name: <Gene name>
~~~

#### Long vs short range TFs

~~~
Use biological knowledge on human genomics to answer the following
classification question that classifies transcriptional factors into
long-range and short-range based on the genomic distances. Answer
with one of these categories: long_range, short_range and respond
with a JSON object containing ‘gene’, ‘answer’, and a short
‘explanation’ without any other supporting commentary or formatting.
Gene name: <Gene name>
~~~

#### NOTCH-1 Targets

~~~
Use biological knowledge on human genomics to answer the following
classification question that classifies N1(NOTCH1) downstream targets from non-targets
factors in the gene network governing cardiac valve
disease. Answer with one of these categories: n1_activated,
n1_nontarget and respond with a JSON object containing ‘gene’,
‘answer’, and a short ‘explanation’ without any other supporting
commentary or formatting. Gene name: <Gene name>
~~~

#### NOTCH-1 Centrality

~~~
Use biological knowledge on human genomics to answer the following
classification question that classifies central versus peripheral
factors within the NOTCH1(N1)-dependent gene network governing
cardiac valve disease. Answer with one of these categories:
n1_central, n1_peripheral and respond with a JSON object containing
‘gene’, ‘answer’, and a short ‘explanation’ without any other
supporting commentary or formatting. Gene name: <Gene name>
~~~

### Small Molecule Property Prediction Prompts

AMES prompts were of the following form, where `<molecule>` is the SMILES string of the small molecule.

~~~
Mutagenicity means the ability of a drug to induce genetic
alterations. Drugs that can cause damage to the DNA can result in
cell death or other severe adverse effects. Nowadays, the most widely
used assay for testing the mutagenicity of compounds is the Ames
experiment which was invented by a professor named Ames. The Ames
test is a short-term bacterial reverse mutation assay detecting a
large number of compounds which can induce genetic damage and
frameshift mutations. Binary classification. Given a drug SMILES
string, predict whether it is mutagenic (1 for active, 0 for
inactive). You must respond with a JSON object, without any
supporting commentary, explanation, markup, or anything else besides
the JSON object. Do not surround the object with backticks or code
formatting. The JSON object should include only two top-level keys:
“answer” and “explanation”, where “answer” contains only a numerical
answer and “explanation” contains only a string with your reasoning
behind the answer. Analyze the following molecule: <molecule>
~~~

#### Bioavailability Ma prompts were of the following form, where <molecule> is the SMILES string of the small molecule

~~~
Oral bioavailability is defined as “the rate and extent to which the
active ingredient or active moiety is absorbed from a drug product
and becomes available at the site of action”. Binary classification.
Given a drug SMILES string, predict the activity of bioavailability
(1 for active, 0 for inactive). You must respond with a JSON object,
without any supporting commentary, explanation, markup, or anything
else besides the JSON object. Do not surround the object with
backticks or code formatting. The JSON object should include only two
top-level keys: “answer” and “explanation”, where “answer” contains
only a numerical answer and “explanation” contains only a string with
your reasoning behind the answer. Analyze the following molecule:
<molecule>
~~~

#### CYP2D6 Inhibition Veith prompts were of the following form, where <molecule> is the SMILES string of the small molecule

~~~
The CYP P450 genes are involved in the formation and breakdown
(metabolism) of various molecules and chemicals within cells.
Specifically, CYP2D6 is primarily expressed in the liver. It is also
highly expressed in areas of the central nervous system, including
the substantia nigra. Binary Classification. Given a drug SMILES
string, predict CYP2D6 inhibition (1 for active, 0 for inactive). You
must respond with a JSON object, without any supporting commentary,
explanation, markup, or anything else besides the JSON object. Do not
surround the object with backticks or code formatting. The JSON
object should include only two top-level keys: “answer” and
“explanation”, where “answer” contains only a numerical answer and
“explanation” contains only a string with your reasoning behind the
answer. Analyze the following molecule: <molecule>
~~~

#### CYP3A4 Substrate Carbon-Mangels prompts were of the following form, where <molecule> is the SMILES string of the small molecule

~~~
CYP3A4 is an important enzyme in the body, mainly found in the liver
and in the intestine. It oxidizes small foreign organic molecules
(xenobiotics), such as toxins or drugs, so that they can be removed
from the body. Binary Classification. Given a drug SMILES string,
predict if it is a substrate to the enzyme (1 for active, 0 for
inactive). You must respond with a JSON object, without any
supporting commentary, explanation, markup, or anything else besides
the JSON object. Do not surround the object with backticks or code
formatting. The JSON object should include only two top-level keys:
“answer” and “explanation”, where “answer” contains only a numerical
answer and “explanation” contains only a string with your reasoning
behind the answer. Analyze the following molecule: <molecule>
~~~

#### CYP3A4 Inhibition Veith prompts were of the following form, where <molecule> is the SMILES string of the small molecule

~~~
The CYP P450 genes are involved in the formation and breakdown
(metabolism) of various molecules and chemicals within cells.
Specifically, CYP3A4 is an important enzyme in the body, mainly found
in the liver and in the intestine. It oxidizes small foreign organic
molecules (xenobiotics), such as toxins or drugs, so that they can be
removed from the body. Binary Classification. Given a drug SMILES
string, predict CYP3A4 inhibition (1 for active, 0 for inactive). You
must respond with a JSON object, without any supporting commentary,
explanation, markup, or anything else besides the JSON object. Do not
surround the object with backticks or code formatting. The JSON
object should include only two top-level keys: “answer” and
“explanation”, where “answer” contains only a numerical answer and
“explanation” contains only a string with your reasoning behind the
answer. Analyze the following molecule: <molecule>
~~~

#### P-gp Broccatelli prompts were of the following form, where <molecule> is the SMILES string of the small molecule

~~~
P-glycoprotein (Pgp) is an ABC transporter protein involved in
intestinal absorption, drug metabolism, and brain penetration, and
its inhibition can seriously alter a drug’s bioavailability and
safety. In addition, inhibitors of Pgp can be used to overcome
multidrug resistance. Binary classification. Given a drug SMILES
string, predict the activity of Pgp inhibition (1 for active, 0 for
inactive). You must respond with a JSON object, without any
supporting commentary, explanation, markup, or anything else besides
the JSON object. Do not surround the object with backticks or code
formatting. The JSON object should include only two top-level keys:
“answer” and “explanation”, where “answer” contains only a numerical
answer and “explanation” contains only a string with your reasoning
behind the answer. Analyze the following molecule: <molecule>
~~~

## Code and Data Availability

All code and data are available here: https://github.com/swansonk14/biomedical_o1

---

**Supplementary Figure 1:**
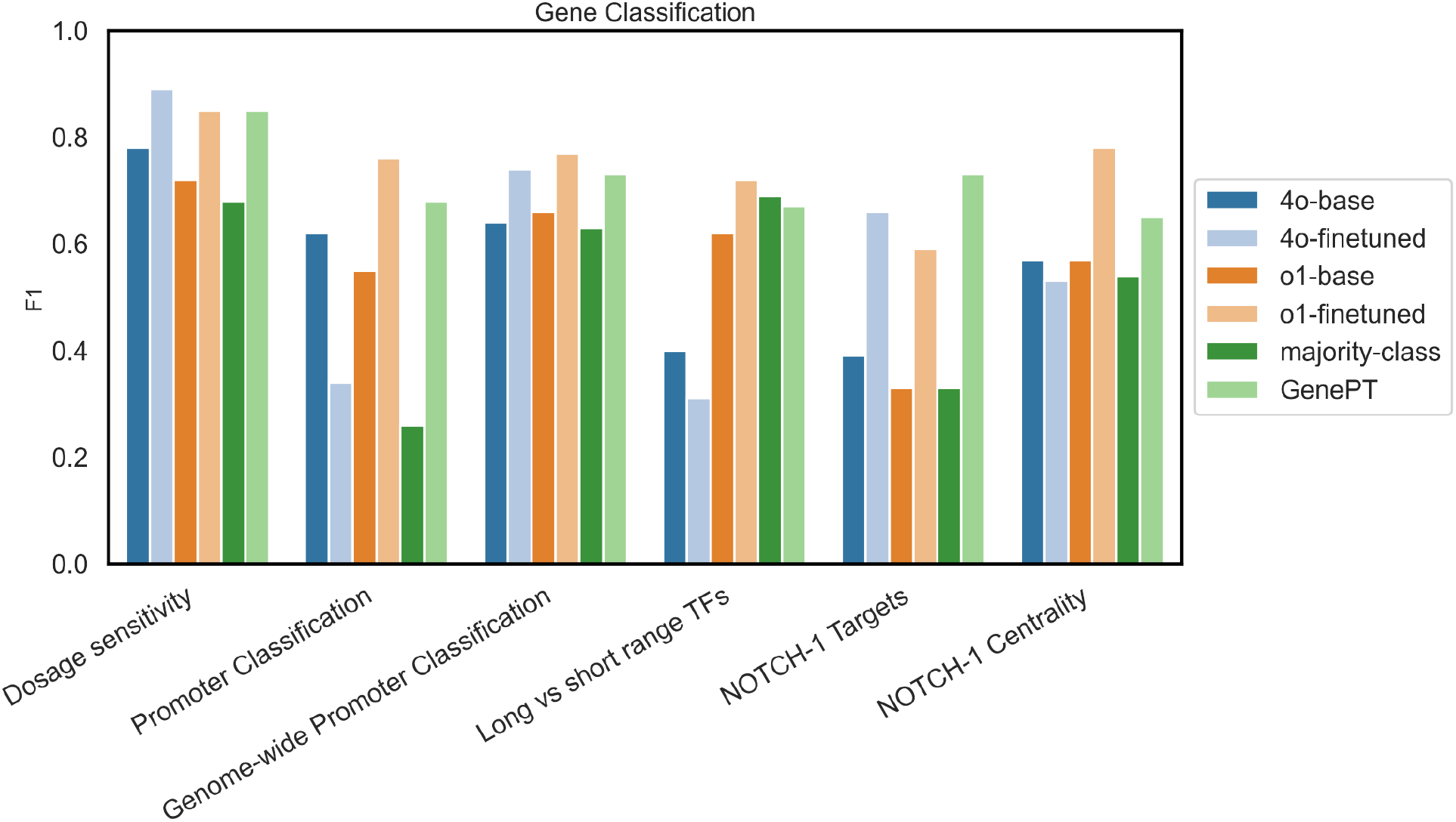
F1 scores (macro-weighted when multi-class) comparing LLM models (before and after finetuning), a majority class baseline, and logistic regression applied to GenePT embeddings on the test sets of gene classification datasets.

**Supplementary Figure 2:**
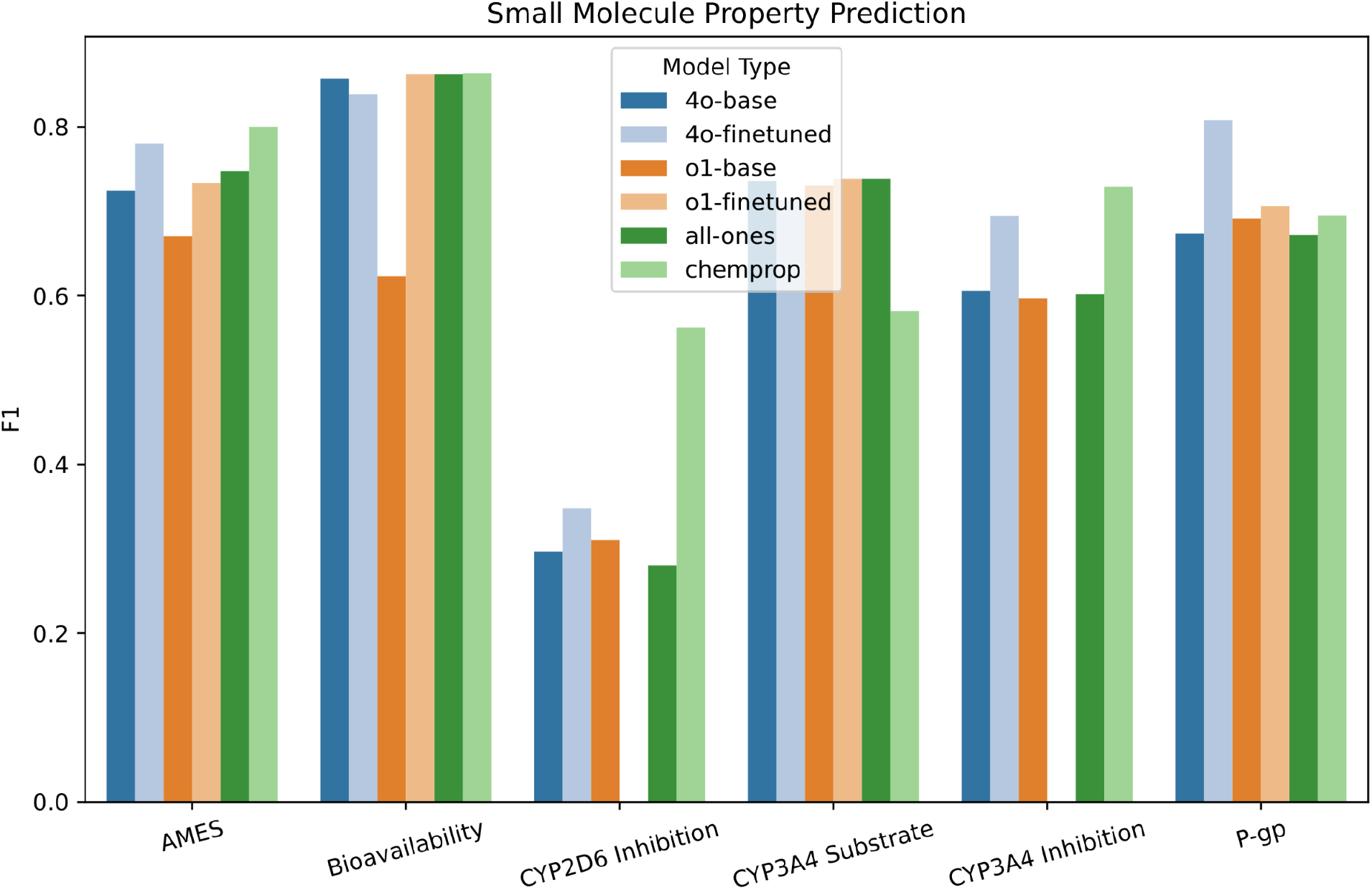
F1 scores comparing LLM models (before and after finetuning), an all-ones baseline, and a Chemprop GNN model on the test sets of the TDC ADMET datasets.

## Appendix

### Antibiotics

In addition to the TDC ADMET small molecule property prediction datasets, we also examined LLM finetuning on an antibiotics dataset. We explored several different data types and LLM training settings, enabling a deeper understanding of the effect of different setups on finetuning performance.

#### Data

The antibiotics dataset (Swanson et al. 2024a) is a dataset of 2,371 small molecules tested for inhibitory activity against the Gram-negative bacterium *Acinetobacter baumannii*. The dataset includes both SMILES and drug names for each molecule, so we experiment with using either SMILES or drug names as input to the LLMs (the Chemprop baseline only uses SMILES). The measurement of inhibitory activity is the normalized optical density at 600 nm (OD_600_), which is a measure of how much bacterial growth occurs after treatment with a molecule. An OD_600_ of 1 indicates that the molecule has no effect since there is equal growth before and after treatment with that molecule. An OD_600_ below 1 indicates inhibition of growth, with smaller values indicating a stronger (better) inhibitory effect. An OD_600_ above 1 indicates increased bacterial growth.

We converted this data to a binary classification problem by creating an OD_600_ threshold of mean minus two standard deviations (OD_600_ = 0.44) across the dataset, with all molecules below this threshold considered active and all molecules above the threshold considered inactive. We used an 80% train, 10% validation, and 10% test split of the molecules using a scaffold split, which ensures that similar molecules end up in the same split—either train, validation, or test—thereby creating a more challenging generalization task from train to test.

Additionally, due to the common phenomenon of mode collapse seen in the TDC ADMET datasets, we experimented with artificially balancing the classes in the train set (the val and test sets were unchanged). We resampled active train molecules to match the number of inactive compounds. In all cases (with or without balancing), if any of the train, validation, or test splits have more than 2,000 molecules, we randomly sample 2,000 molecules from that split (after balancing in the case of balancing).

#### Prompts

Antibiotic prompts were of the following form, where `<molecule>` is the SMILES string or name of the small molecule.

Acinetobacter baumannii is a Gram-negative bacterium that can infect humans. Scientists are interested in developing new drugs that can treat Acinetobacter baumannii infections. Binary classification. Given a molecule, predict whether it can inhibit the growth of Acinetobacter baummannii ATCC 17978 (1 for active, 0 for inactive). You must respond with a JSON object, without any supporting commentary, explanation, markup, or anything else besides the JSON object. Do not surround the object with backticks or code formatting. The JSON object should include only two top-level keys: “answer” and “explanation”, where “answer” contains only a numerical answer and “explanation” contains only a string with your reasoning behind the answer. Analyze the following molecule: <molecule>

#### Experiments

**Appendix Figure 1:**
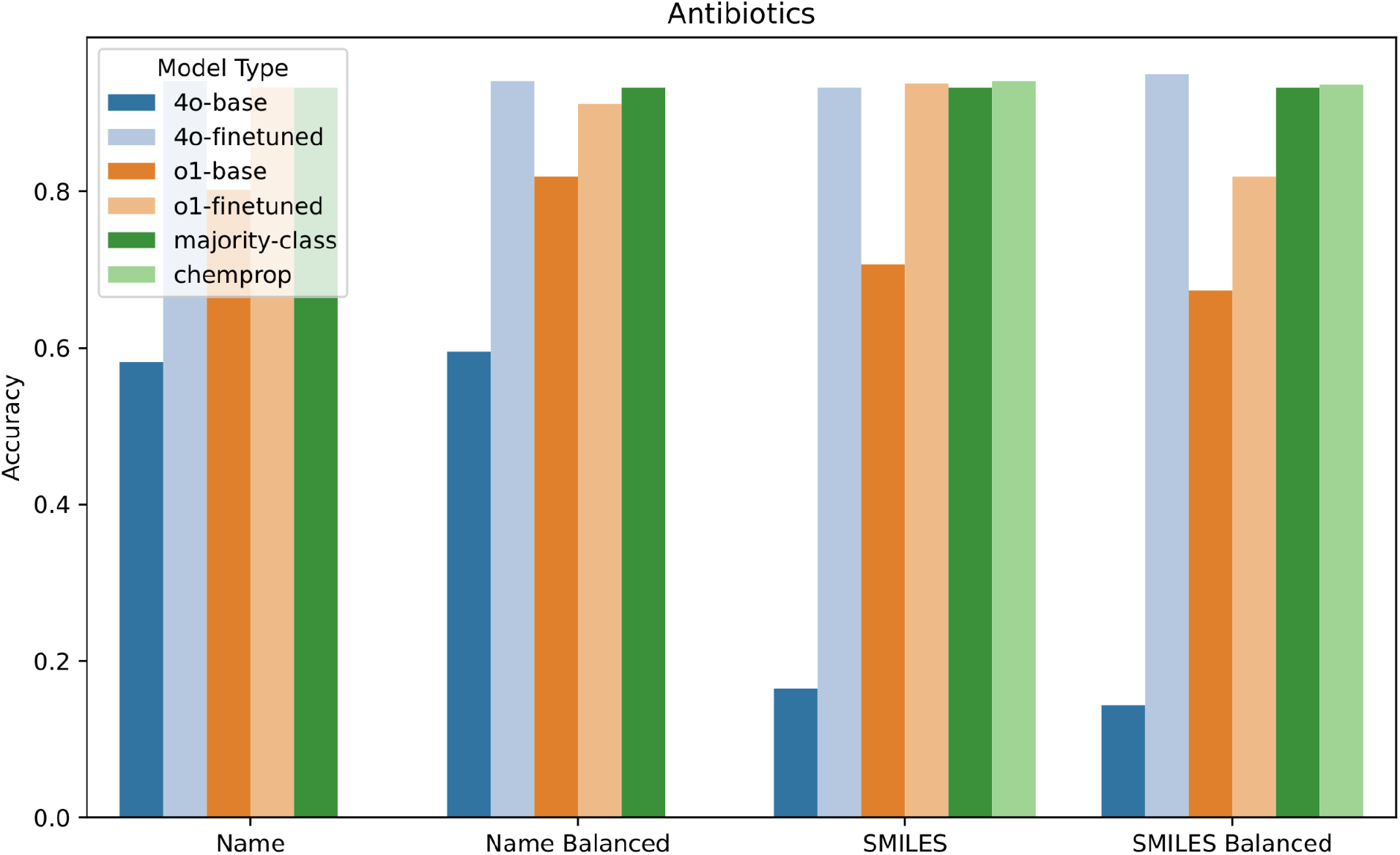
Accuracy scores comparing LLM models (before and after finetuning), a majority class baseline, and a Chemprop GNN model on the test set of the antibiotics dataset, using either SMILES or drug names as input, with and without training set balancing.

**Appendix Figure 2:**
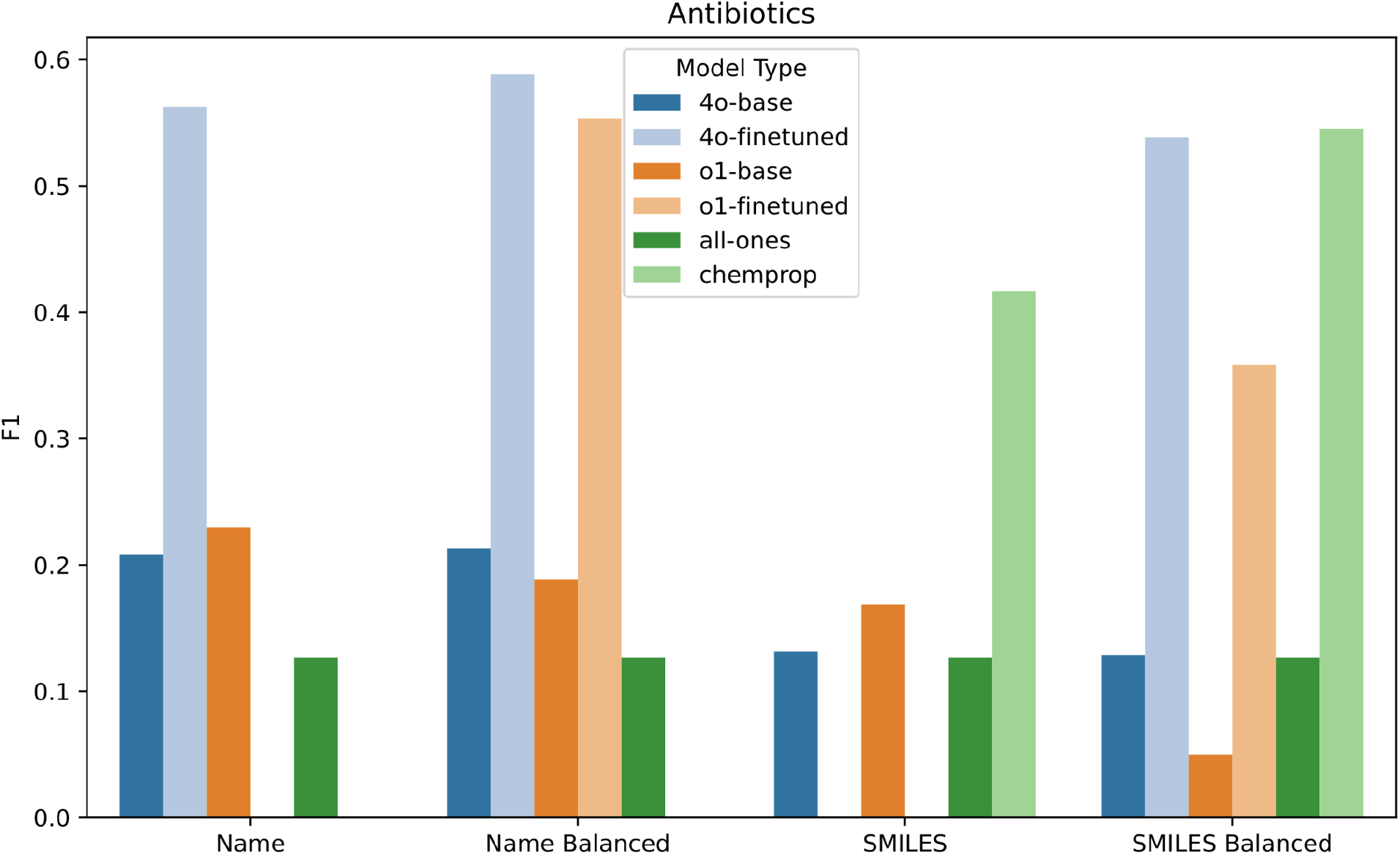
F1 scores comparing LLM models (before and after finetuning), an all-ones baseline, and a Chemprop GNN model on the test set of the antibiotics classification dataset, using either SMILES or drug names as input, with and without training set balancing.

When predicting antibiotic efficacy, the LLMs and Chemprop match the majority class baseline in terms of accuracy (Appendix Figure 1) but significantly exceed the all ones baseline in terms of F1 (Appendix Figure 2), particularly after LLM finetuning. Due to the class imbalance (93% inactive molecules), both 4o-finetuned and o1-finetuned without class balancing often collapse to predicting that all molecules are inactive, which results in high accuracy (equal to the majority class baseline) but an F1 score of 0. However, with class balancing, both 4o-finetuned and o1-finetuned achieve large gains in F1 score relative to 4o-base and o1-base without collapsing to predicting all inactive or all active. Whether using the drug name or the SMILES as input, 4o-finetuned outperforms o1-finetuned. However, neither model is able to outperform Chemprop when using SMILES as input.

## References

Chen, Y., and Zou, J. Y. (2024), “Simple and effective embedding model for single-cell biology built from ChatGPT,” Nature biomedical engineering, Nature Publishing Group, 1–11. 10.1038/s41551-024-01284-6.

Gu, Y., Tinn, R., Cheng, H., Lucas, M., Usuyama, N., Liu, X., Naumann, T., Gao, J., and Poon, H. (2022), “Domain-specific language model pretraining for biomedical natural language processing,” ACM transactions on computing for healthcare, Association for Computing Machinery (ACM), 3, 1–23. 10.1145/3458754.

Heid, E., Greenman, K. P., Chung, Y., Li, S.-C., Graff, D. E., Vermeire, F. H., Wu, H., Green, W. H., and McGill, C. J. (2024), “Chemprop: A machine learning package for chemical property prediction,” Journal of chemical information and modeling, American Chemical Society (ACS), 64, 9–17. 10.1021/acs.jcim.3c01250.

Huang, K., Fu, T., Gao, W., Zhao, Y., Roohani, Y., Leskovec, J., Coley, C. W., Xiao, C., Sun, J., and Zitnik, M. (2021), “Therapeutics Data Commons: Machine learning datasets and tasks for drug discovery and development,” arXiv [cs.LG].

Lee, J., Yoon, W., Kim, S., Kim, D., Kim, S., So, C. H., and Kang, J. (2020), “BioBERT: a pre-trained biomedical language representation model for biomedical text mining,” Bioinformatics (Oxford, England), Oxford University Press (OUP), 36, 1234–1240. 10.1093/bioinformatics/btz682.

Swanson, K., Liu, G., Catacutan, D. B., Arnold, A., Zou, J., and Stokes, J. M. (2024a), “Generative AI for designing and validating easily synthesizable and structurally novel antibiotics,” Nature machine intelligence, Springer Science and Business Media LLC, 6, 338–353. 10.1038/s42256-024-00809-7.

Swanson, K., Walther, P., Leitz, J., Mukherjee, S., Wu, J. C., Shivnaraine, R. V., and Zou, J. (2024b), “ADMET-AI: a machine learning ADMET platform for evaluation of large-scale chemical libraries,” Bioinformatics (Oxford, England), Oxford University Press (OUP), 40, btae416. 10.1093/bioinformatics/btae416.

Theodoris, C. V., Xiao, L., Chopra, A., Chaffin, M. D., Al Sayed, Z. R., Hill, M. C., Mantineo, H., Brydon, E. M., Zeng, Z., Liu, X. S., and Ellinor, P. T. (2023), “Transfer learning enables predictions in network biology,” Nature, 618, 616–624. 10.1038/s41586-023-06139-9.

Wei, J., Wang, X., Schuurmans, D., Bosma, M., Ichter, B., Xia, F., Chi, E., Le, Q., and Zhou, D. (2022), “Chain-of-thought prompting elicits reasoning in large language models,” arXiv [cs.CL].

Yuksekgonul, M., Bianchi, F., Boen, J., Liu, S., Lu, P., Huang, Z., Guestrin, C., and Zou, J. (2025), “Optimizing generative AI by backpropagating language model feedback,” Nature, Nature Publishing Group, 639, 609–616. 10.1038/s41586-025-08661-4.

Zhang, B., Mitchell, E., Ren, H., Lu, K., Schwarzer, M., Pokrass, M., Zhao, S., Sanders, T., Kalai, A., Passos, A., Sokolowsky, B., Le, E. Y., Ritter, E., Sheng, H., Wang, H., Kostrikov, I., Lee, J., Ferstad, J., Lampe, M., Radhakrishnan, P., Fitzgerald, S., Bubeck, S., Dubois, Y., Bai, Y., Applebaum, A., Proehl, E., Mays, E., Parish, J., Liu, K., Maksin, L., Ho, L., Wang, M., Wang, M., Watkins, O., Chao, P., Miserendino, S., Patwardhan, T. A., Woodford, A., Hoover, B., Brill, J., Stirman, K., Ajjarapu, N., Turley, N., Handa, N., Godement, O., Nathan, A., Huang, A., Wang, A., Gohel, A., Eggers, B., Yu, B., Ashley, B., Huang, C., Bogan, D., Sokolova, E., Horacek, E., Such, F., Cohen, J., Gross, J., Becker, J., Wu, K., Lv, L., Byron, L., Liodakis, M., Johnson, M., Trpcic, M., Yesildal, M., Rygaard, R., Marsan, R., Ram-chandani, R., Kshirsagar, R., Conlon, S., Xia, T., Fu, S., Narayanan, S., Choudhry, S., Kaftan, T., Creech, T., Vallone, A., Duberstein, A., Sert, E., Wallace, E., Zhao, G., Kofman, I., Yu, J., Candela, J. Q., Boyd, M.-L., Yatbaz, M., McClay, M., Wang, M., Agarwal, S., Jain, S., Toizer, S., Hernández, S., Mostovoy, S., Li, T., Cha, Y., Wang, Y., Ahmad, L., Peterson, T., Chang, C., Ying, K., Clark, A., Stuckey, D., Tworek, J., Pachocki, J., Heidecke, J.-H., Weil, K., Fedus, L., Chen, M., Altman, S., and Zaremba, W. (n.d.). “OpenAI o3-mini System Card.”

